# Pupil size reveals arousal level dynamics in human sleep

**DOI:** 10.1101/2023.07.19.549720

**Authors:** Manuel Carro-Domínguez, Stephanie Huwiler, Stella Oberlin, Timona Leandra Oesch, Gabriela Badii, Anita Lüthi, Nicole Wenderoth, Sarah Nadine Meissner, Caroline Lustenberger

## Abstract

Recent animal research has revealed the intricate dynamics of arousal levels that are potentially crucial for maintaining proper sleep resilience and memory consolidation. Also in humans, changes in arousal level are believed to be a determining characteristic of healthy and pathological sleep but tracking arousal fluctuations has been methodologically challenging. Here we measured pupil size, an established indicator of arousal levels, during overnight sleep and tested whether the arousal level affects cortical response to auditory stimulation. We show that pupil size dynamics change as a function of sleep macrostructure and microstructural events. In particular, pupil size is inversely related to the occurrence of sleep spindle clusters, a marker of sleep resilience. Additionally, pupil size prior to auditory stimulation influences the evoked response, most notably in delta power, a marker of several restorative and regenerative functions of sleep. Recording pupil size dynamics provides novel insights into the interplay between arousal levels and sleep oscillations, opening new avenues for future research and clinical applications in diagnosing and treating pathological sleep associated with abnormal arousal levels.

## Introduction

Sleep is typically described as a state of low arousal level and disconnection from the environment, characterized by reduced responsiveness to external stimuli. Previous work in rodents has shown that, compared to alert wakefulness, arousal levels are strongly suppressed during non-rapid eye movement (NREM) and rapid eye movement (REM) sleep^1–4^. However, little is known about whether similar arousal level dynamics exist also during human sleep and how they relate to sleep macro- and microarchitecture. To answer this question, we used pupillometry (Fig. 1), a well-accepted method for estimating arousal levels during wake, together with polysomnography (PSG) and electrocardiography (ECG). Recently it has been discovered in rodents that arousal levels exhibit infraslow fluctuations (ISFs) during NREM sleep, representing vigilant and consolidated periods of sleep. The Locus Coeruleus (LC), a small brainstem nucleus and the main source of noradrenergic transmission (NA), seems to be causally mediating these infraslow arousal level fluctuations^1,3^.

**Figure 1.**
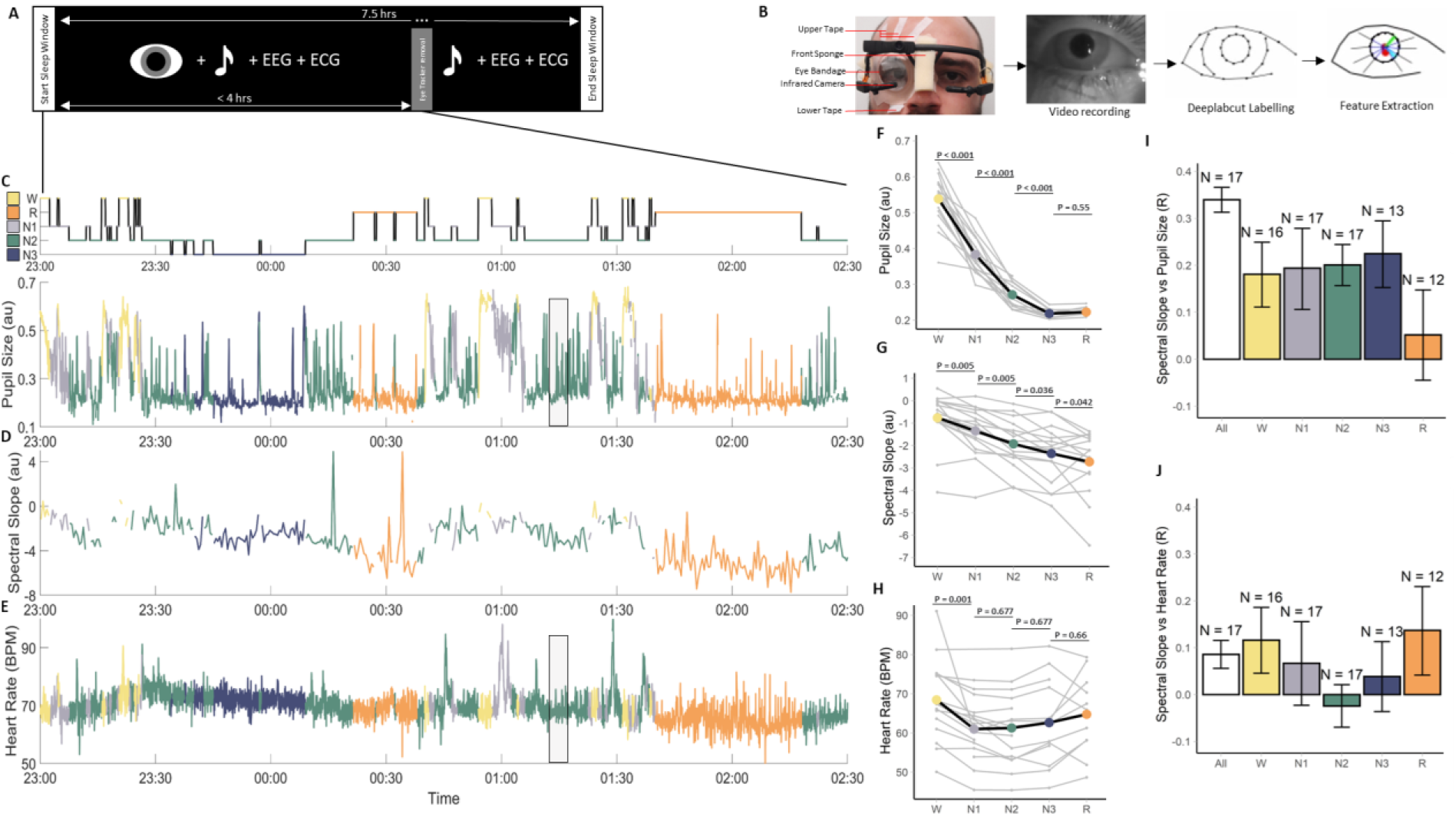
Experimental procedures. (**A**) Overall sleep protocol. Auditory stimulation was applied the whole night during non-rapid eye movement (NREM) sleep whereas pupil size was measured only for up to 4 hours. (**B**) Sleep pupillometry protocol: The eye was taped open and protected with a transparent eye bandage while tracked with an infrared camera. Eye features were extracted from the videos: pupil and eyelid centroids (red markers), fitted ellipse around pupil circumference (black ellipse) major and minor axis of pupil ellipse (cyan and green line), iris points connected to pupil centroid (center black lines). (**C-J**) W: wakefulness (yellow); N1: NREM stage 1 (grey); N2: NREM stage 2 (green); N3: NREM stage 3 (blue); R: rapid eye movement (REM, orange). (**C-E**) Hypnogram while pupil size was recorded and accompanying time series from a representative participant. **(C)** Pupil size fluctuations during sleep. Values are the ratio of the pupil radius to the iris radius. The displayed time period refers to the first 3.5 h of a recording as shown in (A). (**D**) 30-s segments of spectral slope without artifacts. (**E**) Heart rate (HR) in beats per minute (BPM). The grey box is the time period plotted in the bottom plot of Fig. 2A. **(F)** Average pupil size, (**G**) spectral slope, and (**H**) HR for wake, N1, N2, N3, and REM sleep. The data of each participant is depicted in grey. Statistical differences between sleep stages are depicted (see supplementary tables 1-3 for more detailed information) (**I**) Repeated measures correlation within sleep stages where pupil data was aggregated in 30-s segments to match the segment lengths of the spectral slope data. “All” refers to all sleep stages together. N is the number of participants with both valid pupil size and spectral slope for each respective sleep state. (**J**) Repeated measures correlation within sleep stages where HR data was aggregated in 30s segments to match the segment lengths of the spectral slope data. N is the number of participants with both valid pupil size and HR for each respective sleep stage. 30-s segments where auditory stimulation had been applied were excluded.

In mice, ISFs in arousal level occur with a periodicity of approximately 30 seconds to 1 minute^3,5,6^. They show an anti-correlation with spindle activity in the cortex (sigma oscillations in the 12-16 Hz range) and a positive correlation with heart rate (HR). Specifically, consolidated periods of low arousal levels associated with low LC-NA activity coincide with an increased likelihood of sleep spindle clusters, which are believed to protect sleep from external disturbances, and a decrease in HR^3,5–7^. Vigilant periods of high arousal levels associated with high LC-NA activity, by contrast, coincide with low sleep spindle probability and elevated HR. Such periods of elevated arousal levels during NREM sleep might be essential for survival in mice due to the omnipresent environmental threats. Furthermore, recent studies in rodent models indicate that the role of arousal level dynamics during sleep goes beyond changes in vigilance and contributes fundamentally to the macro- and micro-architecture of sleep, coordination of the brain and autonomic system, and memory consolidation potentially playing a fundamental role in restorative processes of sleep (for review, see ^3^).

Yet, it is unclear whether ISFs of arousal levels are preserved in human sleep. Initial evidence indicates that sleep spindles in NREM sleep exhibit an ISF in humans similar to the rodent model^8,9^. However, establishing a systematic link between these spindle fluctuations and changes in arousal level most likely driven by the LC has been challenging due to the lack of appropriate methods for estimating LC activity during human sleep. Here, we propose the use of pupillometry as a novel approach to test several predictions from the rodent literature in human sleep. It is well-established that the size of the pupil’s eye can serve as an indirect marker of arousal levels when light conditions are kept constant^10^. Even though several neuromodulatory systems have been associated with these non-luminance-related changes in pupil size, the LC-NA system has so far received the strongest evidence^11–17^. Furthermore, pupil size and cortical activity have been measured simultaneously during sleep in mice, revealing ISFs of pupil size during NREM sleep that are anti-correlated with electrocortical activity in the spindle range^7^. This finding aligns with previous studies showing LC-NA activity fluctuating with spindle ISFs and indirectly suggests that pupillometry is a valuable tool for tracking arousal levels during sleep and providing a proxy of LC-NA activity during NREM sleep phases^3,5,6,13,18^.

Even though pupillometry is a standard research tool in human participants during wakefulness, very little data has been published reporting human pupil dynamics during sleep^19^. We have therefore developed a safe method for measuring changes in pupil size during sleep using a conventional, low-dose infrared system. Here we synchronized our approach with PSG and ECG. We were able to demonstrate that arousal levels fluctuate in human sleep as revealed by pupil size dynamics across and within sleep stages and established their presence surrounding markers of cortical arousal, sleep electroencephalography (EEG) micro-architectures such as spindles and K-complexes, and HR. Using auditory stimulation during NREM sleep, we further demonstrate that the brain response to administered acoustical stimuli changes depending on arousal levels indicated by differences in pupil size.

## Results

We developed a new approach for measuring pupil size together with PSG and ECG in healthy participants (n=17) during sleep (Fig. 1A-B, Supplementary Video). Portable pupillometry via a low-dose infrared system was applied to the right eye, which was kept open by applying three strips of tape on the upper eyelid and one, wider, strip of tape on the lower eyelid. To reduce discomfort, the eye was taped open without exceeding the subject’s palpebral fissure during wake. To prevent eye dryness, vitamin A-containing eye ointment was applied to the eye immediately before the eye was covered by a transparent eye bandage. Eye bandages are routinely used in nocturnal lagophthalmos patients who cannot close their eyelids during sleep to protect the eye from dust and mechanical damage^20^. All participants could tolerate having their eyes taped open and were able to fall asleep when the room was completely dark except for the infrared light source emitted by the pupillometer. In the debriefing, 2 out of 17 participants reported waking up due to the unfamiliarity of sleeping with one eye open. Participants slept in the lab for 7.5 hours with sleep onset having been adjusted to each individual’s typical bedtime (around 10-11 pm). Auditory stimulation was applied during NREM sleep throughout the night. PSG and ECG were acquired during the whole sleep session but pupillometry was limited to 4 hours, after which the experimenter gently woke the participant up, and removed the eye-tracking equipment. This was a precaution to avoid unwanted side effects such as drying of the right eye. Note that none of our participants reported negative effects, suggesting that longer measurement durations would be achievable. The videos generated by the pupillometer were processed with DeepLabCut^21,22^ to detect pupil size, gaze, and some anatomical features of the eye. Since the eye was taped open without exceeding the subject’s palpebral fissure during wake, there were some invalid video frames when the pupil was occluded by the eyelids. For 15 participants, pupil size could be calculated for at least 44.61% of the video recording (on average, mean±s.e.m., 75.69±5.66% valid samples, n=15). Two participant’s pupils could not be accurately tracked for a large portion of the recording due to head movements resulting in shifting the pupillometer (69.82% invalid samples, n=1) and tape dislodgement resulting in an insufficiently large palpebral fissure (70.37% invalid samples, n=1).

### Pupil size and heart rate reflect dynamic changes in arousal levels at different moments of sleep

Arousal levels can be used to define sleep depth and have been shown to depend on the sleep macro-architecture. To test this basic principle, we determined whether (i) pupil size (Fig. 1C), an established indicator of central arousal levels during wake (ii) EEG spectral slope (Fig. 1D), a marker of cortical arousal^23^, and (iii) HR (Fig. 1D), a marker for cardiovascular activation, differed across sleep stages using linear mixed effect models and calculating post-hoc t-tests corrected for multiple comparisons using Benjamini-Hochberg correction. Pupil size, measured as the ratio between the pupil radius and the iris radius, was largest when being awake and significantly decreased for NREM stage 1 (N1), NREM stage 2 (N2), NREM stage 3 (N3), and REM sleep (Fig. 1F,F(4, 57.43)=168.90, p<0.001). The strongest changes were observed from wake to N1 and from N2 to N3, while pupil size was consistently small for N3 and REM sleep (Supplementary Table 1). These results show that pupil changes across sleep stages in humans and are consistent with previous pupillometry findings in sleeping rodents^7,24^.

To assess whether pupil size can index cortical arousal, we compared pupil size to spectral slope, an electrophysiological marker proposed to reflect the synaptic excitation to inhibition balance with more negative spectral slopes indicating enhanced inhibition^25^. The spectral slope was calculated in 30-s segments in the 30-45Hz range of the Fpz electrode^23^. Consistent with previous studies^23,26,27^, we observed a significant decrease from wake to REM sleep in a near-linear fashion (Fig. 1G, F(4, 68)=41.37, p<0.001, Supplementary Table 2). Repeated measures correlation analyses revealed a significant positive correlation between pupil size and spectral slope across (R=0.34, p<0.001) and within sleep stages (Fig. 1I, all R>=0.18, all p<0.001), except for REM sleep (R=0.05, p=0.291).

HR showed a U-shaped pattern from wake, through NREM sleep stages, to REM sleep (F(4, 55.52)=7.623, p<0.001), whereby only HR in NREM sleep stages was significantly lower compared to wake (Fig. 1E, Supplementary Table 3). Relating HR and spectral slope, there was a weak yet significant correlation across sleep stages (R=0.09, 95%CI=[0.056, 0.116], p<0.001, N=17). Moreover, HR was significantly but weakly related to spectral slope for wake (R=0.12, p=0.001) and REM (R=0.14, p=0.005), but not for NREM sleep stages (Fig. 1J, R≤0.07, p≥0.143).

In summary, our findings suggest that pupil size changes across sleep stages and appears to be linked to cortical arousal. HR also varied across sleep stages but seems to reflect the overall cardiovascular activation stage which is only weakly associated with cortical arousal markers.

### Distinct cortical and cardiovascular states are present across the arousal level spectrum of N2 sleep

Spindle trains have been shown to exhibit infraslow fluctuations (ISFs) in rodents and humans^8,9,28^. Similar ISFs that are phase-locked to spindle clusters occur in the HR and the LC-mediated thalamic noradrenergic levels, where causal evidence shows that LC activity needs to be low for spindle trains to occur^5,6^. To test whether spindle ISFs also exist in humans we calculated Cz sigma power (spindle frequency range of 12-16Hz) during N2 (see Supplementary Fig. 1A for verifying the spindle ISF detection algorithm). We further tested whether similar infraslow dynamics are also observed for pupil size, HR, and K-complexes, a key EEG feature of human N2 sleep that has been argued to reflect arousal level fluctuations that are hallmarked by widely synchronized changes in excitability^29^. Figure 2A-C shows exemplary recordings of sigma power fluctuations (representing spindle ISF, black lines) together with pupil size (Fig. 2A), HR (Fig. 2B), and K-complexes (Fig. 2C, green dots) during N2 sleep. It can be seen that all signals exhibited oscillations on infraslow timescales. Next, we applied a cycle-by-cycle analysis to sigma power (see Methods for details) and identified four phases marking sigma power’s rise, peak, fall, and trough (Fig. 2D-F, black line). We then plotted pupil size (Fig. 2D), HR (Fig. 2E), and K-complex likelihood (Fig. 2F) as a function of the total spindle ISF cycle length (in %). All parameters exhibited significant modulations as confirmed by statistically comparing average pupil size, HR, and K-complex likelihood between the rise, peak, fall, and trough phase of the sigma power cycle using linear mixed effect models (all F(3, 48) >= 10.85, all p< 0.001, Fig. 2G-I, Supplementary Tables 4-6).

**Figure 2.**
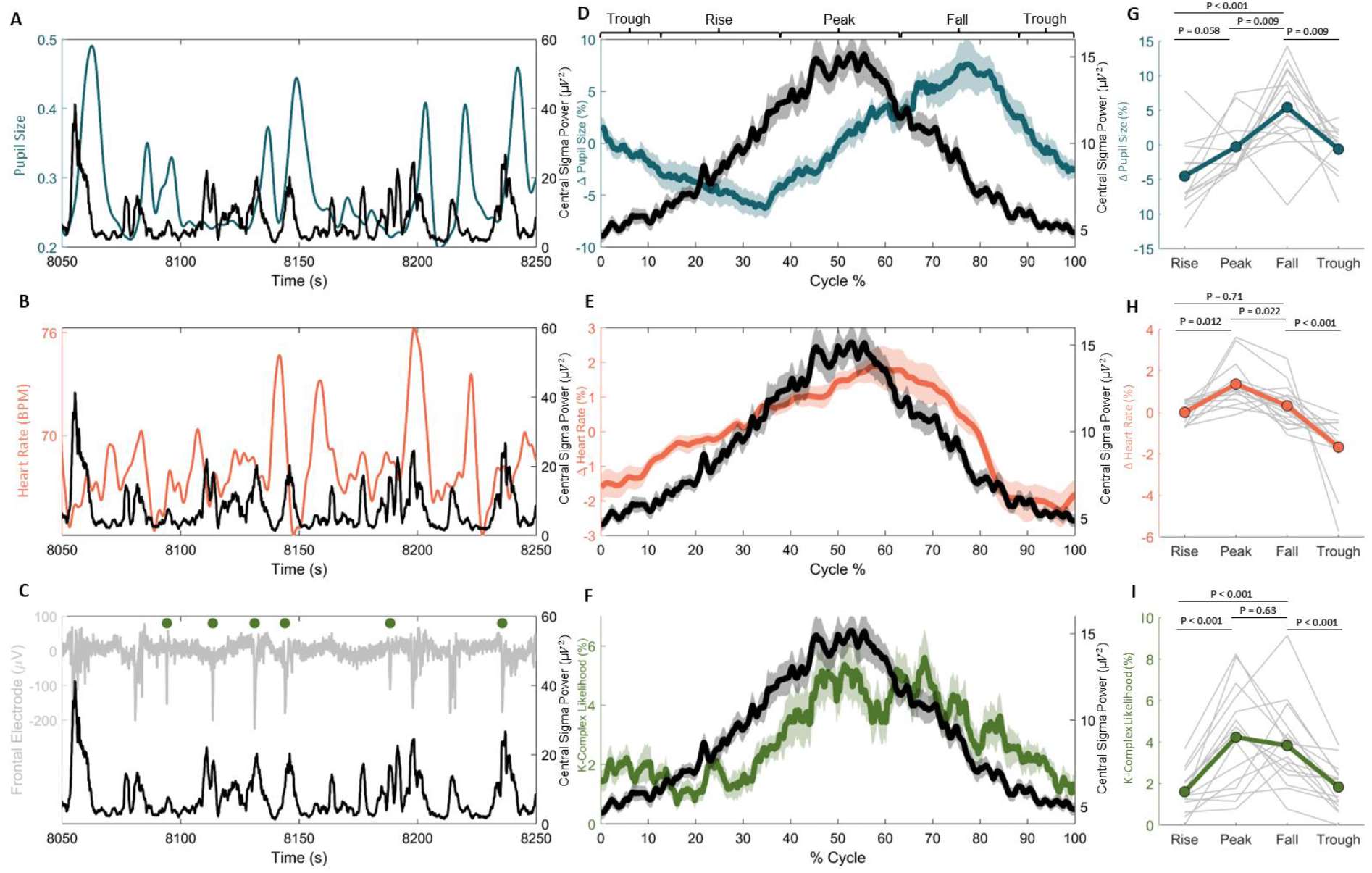
Relationship between sigma power, pupil size, heart rate, and K-complexes. (**A-C**): Individual example of sigma power fluctuations alongside the corresponding pupil size, HR, and K-complexes (green dots in C), respectively. The example window corresponds to the time in the grey boxes in the sleep recording from Fig. 1C and 1E. (**D-F**) Sigma power (black & right axis) with pupil size (blue & left axis in **D**), heart rate (red & left axis in **E**), and K-complex likelihood (green & left axis in **F**). All metrics are normalized to the timing of the troughs of detected spindle infraslow fluctuations (ISF). The time between troughs was renormalized in terms of cycle percentage. Centre line plots represent the mean across participants and shadings around center lines represent the standard error of the mean (SEM). (**G**) Average pupil size, (**H**) heart rate, and, (**I**) K-complex likelihood for the rise, peak, fall, and trough of the normalized sigma power fluctuations. The data of each participant is depicted in grey. Statistical differences between the rise, peak, fall, and trough of sigma power ISFs are depicted, p-values are based on post-hoc t-test and are adjusted for multiple comparisons using Benjamini-Hochberg correction.

Sigma power and pupil diameter appear to oscillate in a slightly phase-shifted manner (Fig. 2D). Cross-correlating pupil size and sigma power (Supplementary Fig. 1B) revealed that the two signals are shifted by approximately 30% of the cycle (corresponding to a 108-degree phase difference with R=-0.26). Pupil size is smallest during the rise of sigma power and reaches its peak when sigma power is falling again (Fig. 2G). Note that pupil size was significantly larger during the fall of sigma power than during any other phase (p<=0.009).

Contrary to pupil diameter, HR and K-complex likelihood fluctuations tended to occur phase-locked to sigma power fluctuations (Fig. 2E, F). A cross-correlation between sigma power and HR revealed a delay of the HR fluctuation of approximately 5% (corresponding to an 18-degree phase difference, R=0.26) and HR was significantly higher during the peak phase of sigma power than during the rise or fall (p<=0.022). K-complexes were significantly less likely to occur during the rise of sigma power than during the peak and fall of sigma power (both p<0.001).

In summary, we show for the first time in humans that fluctuations in pupil size, HR, and K-complex likelihood are linked to spindle ISFs. Spindle ISFs are an EEG marker that has been previously shown to be linked to LC-mediated arousal levels, i.e. spindle occurrence is low when LC-NA levels are high and vice versa during NREM sleep in rodents^5,6^.

### Pupil size and heart rate fluctuations reveal distinctly different dynamics across N2 events

Next, we investigated how pupil size and heart rate relate to three specific events in N2 sleep micro-architecture that hallmark changes in arousal levels, i.e. K-complexes, spindle clusters, and sleep arousals as defined in the American Academy of Sleep Medicine (AASM) criteria^8,30–33^. Although sigma power provides an indirect but continuous readout of spindle activity, discretizing the Cz signal into spindle clusters provides a uniform timescale in addition to a more sensitive and precise readout of strong spindle activity. It is worth noting that not all participants exhibited each event type together with valid pupil size or HR measurements during epochs with unperturbed sleep (Fig. 3). To evaluate if pupil size and HR are different in relation to these sleep-microarchitecture events and therefore distinct from the overall N2 state, we investigated pupil and HR dynamics at each N2 event type and used t-tests corrected for multiple comparisons using Bejamin-Hochberg correction.

**Figure 3.**
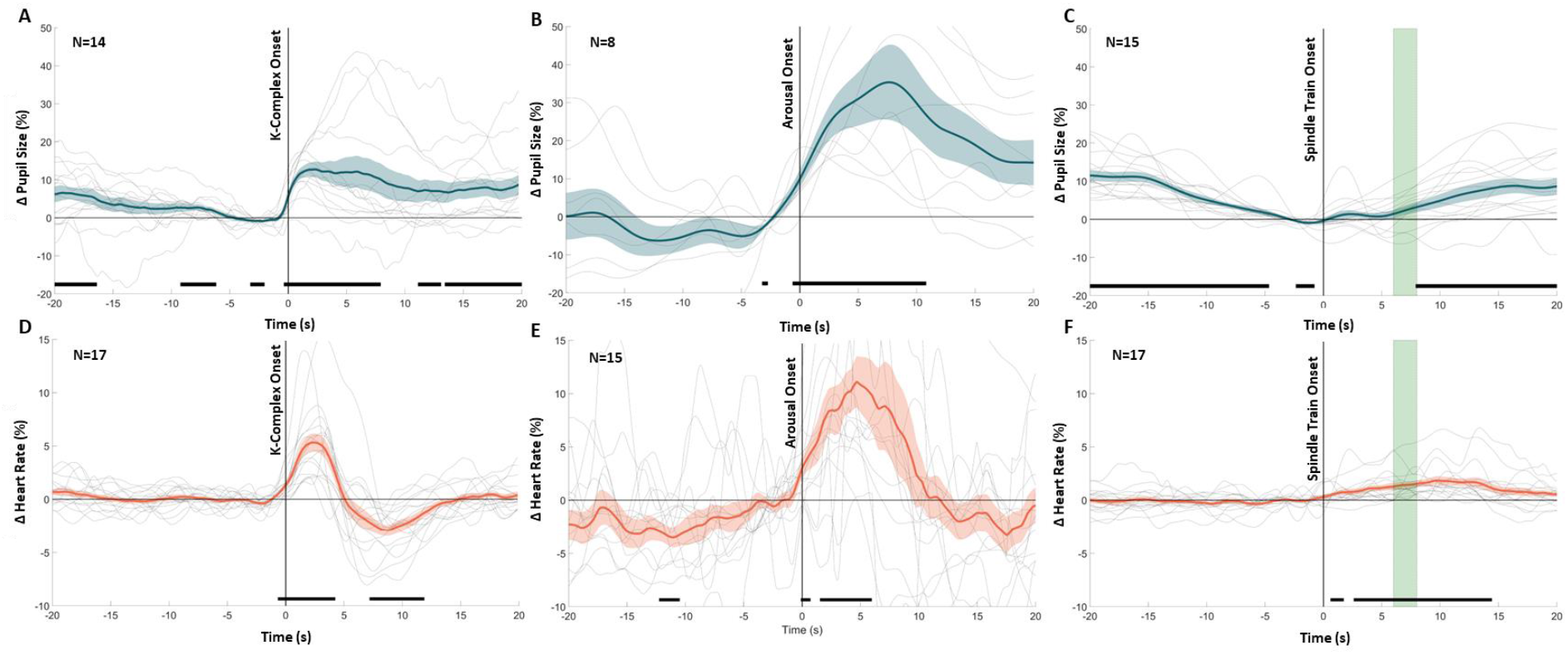
Pupil size and instantaneous heart rate dynamics of N2 events. (**A-C**) Pupil size color-coded in blue and normalized to the 5s prior the detected event: K-complexes, arousals, and spindle trains (respectively). (**D-F**) Heart rate color-coded in orange and normalized to the 5s prior to the detected event: K-Complexes, spindle trains, and arousals (respectively). (**C, F**) Green vertical shading in top panels reflects the mean ± 1 SEM of the median spindle train end across participants. Centre lines represent the group mean and shading around the centre lines the SEM. Grey lines are the mean response of each participant. Black horizontal lines mark significant differences from the 5s baseline prior to each event.

For K-complexes (Fig. 3A), we observed a clear pupil dilation within the 10s after the K-complex with an average increase of 12.84±2.12% from baseline. The black bars in Fig. 3A represent regions significantly different from baseline. Interestingly, systematic changes in pupil diameter are revealed that appear to precede the K-complex onset by approximately 1s (p<0.05) and last up to approximately 4s (p<0.05) after the onset. Figure 3A also shows that average pupil size gradually decreased to a minimal level before the dilation started. Furthermore, HR revealed a biphasic response to K-complexes reflected in an initial increase of up to 5.34±0.85% from baseline (p<0.05) followed by a decrease of up to 2.96±0.48% below baseline (p<0.05, Fig. 3D). This biphasic HR response to K-complexes aligns with previous findings^32^.

Arousals (Fig. 3B) from sleep as defined by AASM guidelines are brief perturbations to sleep which are characterized by large and sudden changes in the theta, alpha, or beta frequencies in the EEG power spectra, which can be accompanied by EMG activation. We found sleep arousals to be accompanied by a pupil dilation that typically started slightly before the determined AASM arousal onset and reached a significant increase of up to 35.37±9.88% compared to baseline, around 7s after AASM arousal onset (p<0.05). This pupil dilation was descriptively larger than the dilations observed during K-complexes but also more variable across participants. When comparing pupil size dynamics between K-complexes and arousals, statistics did not survive correction for multiple comparisons. Similarly, HR increased by 11.18±2.31% in parallel with arousals (p<0.05, Fig. 3E) which is descriptively but not statistically larger in magnitude than HR variations observed during K-complexes (Fig. 2D-E).

Spindles (Fig. 3C) and their clustering into “trains” occur during low LC-NA activity (low arousal level) and have been argued to play a sleep-protecting role^30,34^. In line with the pupil size fluctuations during spindle ISFs (Fig. 2D, G), pupil size exhibited a U-shaped pattern during the 40-s period surrounding spindle trains. Prior to the onset of the spindle train, there was a significant 11.51±1.46% decrease in pupil size compared to baseline (p<0.05). Pupil size remained relatively small throughout the spindle train and only after did it return to values observed 20s prior to the onset of the spindle train (p<0.05). We found that HR started to slowly increase at the onset of spindle trains and remained 1.851±0.42% higher than baseline beyond the end of most spindle trains (p<=0.05, Fig. 2F).

To shed light on the pupil size and HR dynamics surrounding these three events, we compared the baseline of each event to the average pupil size (F(3, 39.30)=3.58, p=0.022) and heart rate (F(3, 40.027)=6.11, p=0.002) during periods of N2 that were event-free. Pupil size tended to be lower, although not significantly, than general N2 levels prior to K-complexes (-8.30±2.31%, p=0.071) and spindle trains (-8.34±2.75%, p=0.071) but not arousals (0.95±4.94%, p=0.717). Conversely, heart rate was significantly higher prior to arousals (3.50±1.59%, p=0.008) but was not significantly different from general N2 levels prior to K-complexes (-1.03±0.46%, p=0.382) and spindle trains (-0.89±0.76%, p=0.382, Supplementary Table 7).

In summary, the pupil constricted before K-complexes and spindle trains, with HR remaining stable, and dilated following all three events while HR showed less pronounced responses to these events (Supplementary Fig. 2).

### Cortical response to sensory stimulation varies with arousal levels

Finally, we investigated whether arousal levels as estimated from pupil size affect cortical response to external stimuli. We applied auditory stimulation (verum) at varying arousal levels, and a control condition where no stimuli were applied (sham, Fig. 4). To statistically compare verum versus sham conditions, we averaged the data within three time bins (i) Stim_early_ (first 5s of the auditory stimulation), (ii) Stim_late_ (last 5s of the auditory stimulation), (iii) Stim_post_ (5s bin after auditory stimulation was switched off) and applied paired t-tests between verum and sham for each time bin and corrected for multiple comparisons using Benjamin-Hochberg correction (Fig. 4; Methods). We found that the pupil showed a sharp dilation during Stim_early_ and Stim_late_ that was significantly larger for verum than for sham stimulation (Fig. 4A, D). This dilation reached its peak shortly after the end of the 10s stimulation window upon which pupil size gradually decreased again. Sigma power, being associated with an increased likelihood of sleep spindle occurrence, significantly increased during verum Stim_early_ when compared to sham, which is typically observed when slow wave amplitude is increased by tones^35,36^. However, after the initial peak, sigma power decreased quickly and was significantly suppressed during Stim_post_ relative to sham (Fig. 4B, E). In parallel, delta power representing slow wave activity increased during Stim_early_ and Stim_late_ when compared to sham, and then during Stim_post_ returned to Stim_pre_ values (Fig. 4C, F).

**Figure 4.**
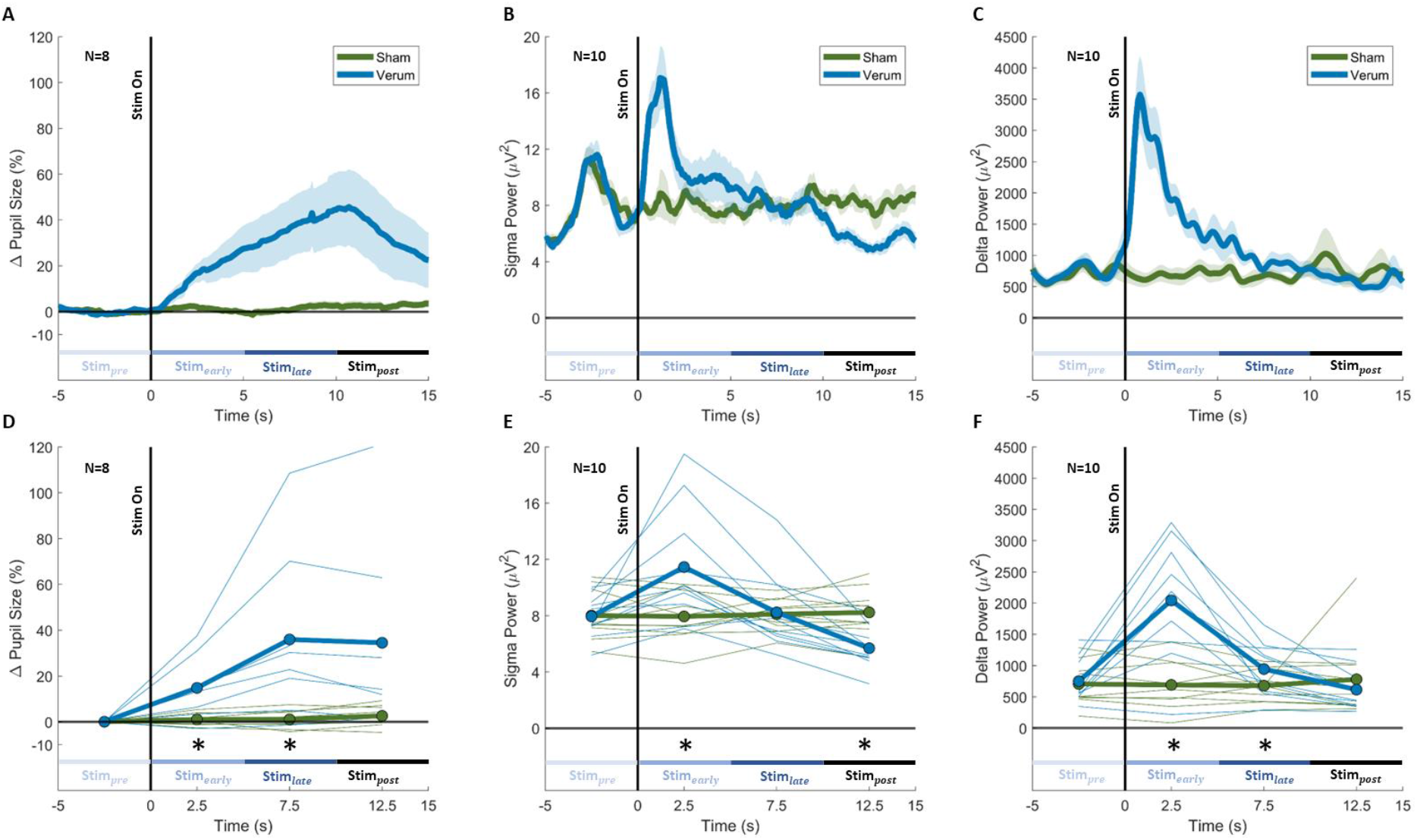
Pupil and EEG responses to auditory stimulation. We applied a 10-s stimulation condition of 1 Hz rhythmic auditory stimulation at 45dB (verum) at varying arousal levels and a control condition where no stimuli were applied (sham). (**A**-**C**) Pupil size, sigma power, and delta power relative to the 5s prior to the detection of verum (blue) and sham (green) stimulation windows. Centre lines represent the mean and shading around centre lines represents SEM. Time zero is the detection of the stimulation window. (**D**-**F**) Pupil size, sigma power, and delta power of **A**-**C**, respectively, during verum (blue) and sham (green) averaged every 5s from time zero onwards for each participant. Black stars highlight significant differences (p<0.05) between verum and sham for a given 5s region. (**D**) Pupil in verum condition was significantly larger than sham during Stim_early_ (13.77±4.81%, p=0.046) and Stim_late_ (34.87±12.90%, p=0.046). (**E**) Sigma power in verum condition was significantly larger during Stim_early_ relative to sham (3.52±1.30μV^2^, p=0.036) and significantly lower during Stim_post_ relative to sham (-2.55±0.46 μV^2^, p=0.001). (**F**) Delta power in verum condition increased during Stim_early_ (1’353±271.54μV^2^, p=0.002) and Stim_late_ (265.95±78.29μV^2^, p=0.001) when compared to sham and then during Stim_post_ returned then back to Stim_pre_ values (-165.23±135.99μV^2^, p=0.255).

We therefore investigated whether pupil size, as a potential indirect proxy of central arousal levels, prior to playing a tone modulates the response of sigma and delta power to auditory stimulation. Therefore, we separated N2 verum trials where pupil size during Stim_pre_ was large (highest quartile, representing “high” arousal level) versus small (lowest quartile, representing “low” arousal level) and compared the sigma and delta power 5s prior to stimulation onset (Stim_pre_) with the first 5s of stimulation (Stim_early_) (Fig. 5A-B; Methods). Linear mixed effect models revealed there was a significant difference between large and small pupil size during Stim_pre_ to general N2 levels (F(2, 14.00)=47.06, p< 0.001). Post-hoc t-tests corrected for multiple comparisons revealed that large pupil size during Stim_pre_ was greater than general N2 levels (18.54±5.50%, p=0.004) and small pupil size baselines were smaller than general N2 levels (-25.21±3.40%, p<0.001). There was an increase in sigma power from Stim_pre_ to Stim_early_ in the small pupil baseline condition but not in the large pupil baseline condition (Fig. 5A). Furthermore, there was an increase in delta power from Stim_pre_ to Stim_early_ in the small pupil baseline conditions (928.67± 381.79μV^2^) but not in the large pupil baseline conditions (-105.90± 277.54μV^2^, Fig. 5B).

**Figure 5.**
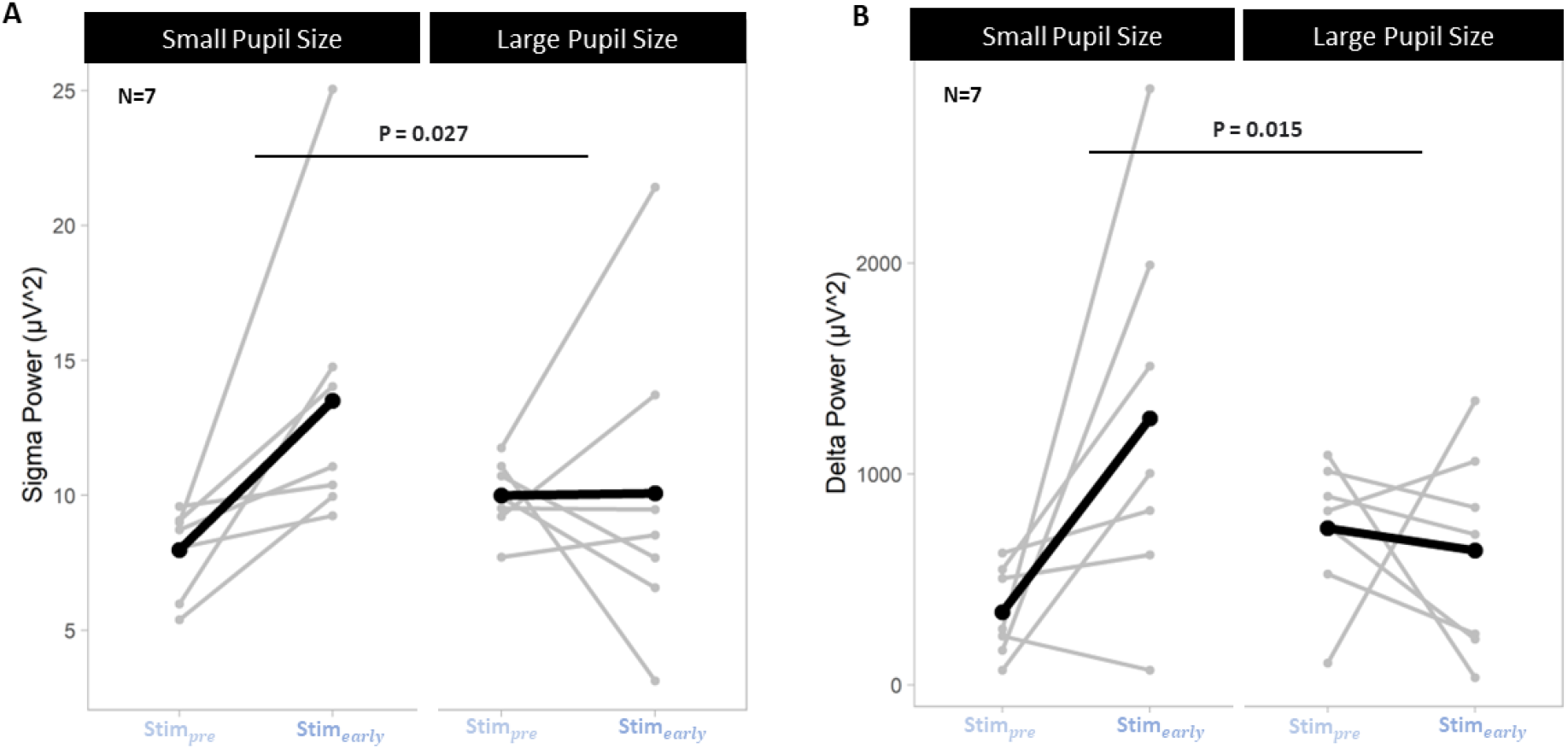
EEG responses to auditory stimulation at small and large pupil size baselines. (**A**) Sigma power 5s before (Stim_pre_) and during the first 5s (Stim_early_) of verum stimulation of trials with large (highest quartile) and small pupil baselines (lowest quartile) to group into high and low arousal level trials. We found an interaction effect indicating increased sigma power from Stim_pre_ to Stim_early_ in the small pupil baseline condition (5.53±2.04μV^2^) but not in the large pupil baseline condition (0.08±2.17μV^2^, F_sigmaXpupilrange_(1, 21)=5.66, p=0.027, post-hoc comparison Stim_pre_ to Stim_early_ with small pupil baseline: F(1, 7)=8.60, p=0.044, post-hoc comparison Stim_pre_ to Stim_early_ with large pupil baseline: F(1, 7)=0.002, p=0.97. (**B**) Delta power during Stim_pre_ and during Stim_early_ of verum stimulation of trials with large (highest quartile) and small pupil baselines (lowest quartile) to group into high and low arousal level trials. We found an interaction effect indicating increased delta power from Stim_pre_ to Stim_early_ in the small pupil baseline condition but not in the large pupil baseline condition F_deltaXpupilrange_(1, 28)=6.77, p=0.015, post-hoc comparison Stim_pre_ to Stim_early_ with small pupil baseline: F(1, 14)=7.58, p=0.031, post-hoc comparison Stim_pre_ to Stim_early_ with large pupil baseline: F(1, 14)=0.25, p=0.623. The horizontal black lines connecting pupil size ranges are the interaction effects between pupil size range and Stim_pre_-Stim_early_ power values.

In summary, our findings show for the first time that auditory stimulation during sleep induces a pupil response upon stimulation. Furthermore, we demonstrate distinct cortical responses to auditory stimulation at high and low arousal baseline levels as indicated by pupil size prior to stimulation onset.

## Discussion

In the present study, we monitored pupil size during human sleep using a low-dose portable infrared system and compared it to cortical and cardiovascular events measured via polysomnography and ECG. We demonstrate that pupil size dynamics relate to macro- and microarchitecture of sleep and predict the brain’s response to auditory stimuli. Therefore, we propose that pupil size dynamics might represent a non-invasive way to track arousal levels.

We showed in humans that pupil size decreases significantly from wakefulness to deep NREM sleep, reaching an equally small diameter in N3 and REM sleep. Similar pupil size changes across sleep states have previously been reported in rodent models^7^. Here, we reproduced the pronounced decrease from wakefulness to NREM sleep (N1 and N2) and then to N3 and REM sleep. Note that there is no distinction of NREM sleep stages in rodents so that the small pupil size which characterizes not only REM sleep but also NREM N3 sleep has remained undetected until now.

In the present study, observed changes in pupil size were paralleled by a decrease in spectral slope, an electrophysiological marker of cortical arousal level^23^, as NREM sleep deepened. Spectral slope levels decreased nearly linearly across NREM sleep stages. These findings suggest that pupil dynamics capture pronounced changes in vigilance state. Unlike pupil dynamics, spectral slope did further decrease from N3 to REM sleep, suggesting that spectral slope and pupil diameter might reflect slightly different physiological mechanisms during REM sleep even though it cannot be ruled out that further pupil constriction was prevented due to a floor effect. HR, on the other hand, showed little to no association with the spectral slope, across sleep stages and also within NREM and wake, indicating that this measure seems to be less closely linked to cortical arousal level dynamics.

During the infraslow fluctuations of sigma power in N2, we found pupil size, a marker of arousal levels, to be low before high sigma (12-16 Hz) power, which reflects high sleep spindle probability (Fig. 2A, D, G, Supplementary Fig. 1B). Similarly, pupil size measured in rodents was found to be low during high sigma power, which also reflects high sleep spindle probability in mice^7,8^. Furthermore, this is also in line with recent studies in rodents, which have demonstrated that arousal levels mediated by LC-NA activity exhibit pronounced infraslow fluctuations (ISFs) within NREM sleep which negatively correlate with the occurrence of sleep spindles. Interestingly, there is causal evidence showing that experimentally increasing arousal levels suppresses the occurrence of sleep spindles which also causes elevated response to sensory stimuli in thalamocortical and autonomic circuits. Conversely, decreasing arousal levels enables spindle occurrence which has been shown to be associated with suppressed response to sensory stimuli^5,6,8,13,30^. Drawing from our results and previous research in rodents, pupil size might be associated with LC-mediated arousal levels during human NREM sleep. However, as discussed below, other brain regions are likely to be involved in the regulation of arousal level and pupil size.

During the infraslow fluctuations of sigma power in N2, we found HR, a marker commonly used to infer autonomic activity during sleep, to be high while sigma power was high (Fig. 2B, E, H, Supplementary Fig. 1C). This is in line with research in humans but, interestingly, not with mice where it was found that heart rate was low during high sigma power^7,8^. Lecci et al. suggested that this dissimilar phase relationship between HR and sigma power may be related to anatomical and physiological differences between the neural coupling to the heart in both species. Although our findings and previous literature suggest a tight coupling between heart rate and central arousal levels at an infraslow timescale (Supplementary Fig. 1B, C), there are no studies to our knowledge that have investigated central arousal levels or HR fluctuations during individual spindle trains, a more sensitive and temporally precise approach to capture spindle activity underlying sigma power (Supplementary Fig. 1A)^37^. In general, if we want to fully understand the link between autonomic activity and cortical arousal during fluctuating arousal levels, we may need to look at sleep microstructural events at a smaller timescale.

We therefore established how pupil dynamics evolve around such microstructural events and observed changes in pupil size prior to, during, and after the start of K-complexes, spindle clusters, and sleep arousals. More specifically, we found that pupil size was smallest right before the onset of the K-complex followed by pupil dilation. This pupil dynamic is reminiscent of that reported in animal literature where arousal levels were found to increase during short pupil constrictions and subsequent dilations^17,38,39^. Pupil size constricted before spindle clusters, perhaps as a result of the reduced arousal levels suggested to be required for spindles to occur^5,39^. Therefore, a reduction in arousal levels seems to be tightly linked to the likelihood for sleep-related cortical events such as K-complexes and sleep spindles to occur, a relationship that can be captured by measuring pupil size. Conversely, HR remained largely unaffected prior to the occurrence of sleep spindles and K-complexes, but showed an increase following the onset of these events. Therefore, HR seems to be less closely linked to cortical arousal level dynamics prior to an arousal event per se but may rather be used to classify the autonomic response to such events^31^. In rodents, Eschenko et al. reported LC firing to be time-locked to specific phases of slow waves^40^. Here, we specifically focused on K-complexes that represent highly synchronized bottom-up slow waves that potentially involve LC activation and the promotion of highly synchronized cortical activity^41,42^. Along these lines, Osorio-Forero et al. (2023) reported that transient LC activity increased delta power, possibly reflecting K-complexes^43^. Our results indicate that K-complexes and sleep arousals both involve a significant pupil dilation possibly reflecting transient LC activity time locked to these events.

Assuming that different pupil levels track different arousal levels, we would expect that the cortical response to auditory stimulation differs depending on when these stimuli are played as a function of pupil size. According to previous research, we found auditory stimulation to increase delta and sigma power dynamics^35,44–50^. These delta responses are believed to involve enhanced slow waves or K-complexes^51,52^. However, this increase in delta power was indeed only present when pupil size was low (lowest quartile) prior to stimulation onset. Therefore, low pupil size might be a prerequisite for the induction of sensory-evoked delta enhancement. It has been previously proposed that K-complexes can only be evoked during times of lower blood pressure^53^ and lower HR^32^, indicating decreased cardiovascular activation, or likely a state of decreased arousal. In rodents, Hayat et al. (2020) showed that tonic LC activity levels were higher before tones that led to sensory-evoked NREM sleep perturbations compared to tones resulting in maintenance of NREM sleep^13^. Importantly, in both cases the tones induced a phasic firing of LC time-locked to the tone onset, but only high prior arousal levels resulted in sleep arousals. In this regard, our results might point to the idea that during low arousal levels, as indicated by low pupil size here, slow oscillations might be induced as a consequence of the short phasic firing of LC which may act sleep protective. However, our stimulation protocol did only involve low volume sounds that are generally not inducing sleep perturbations. Overall, our results indicate that the arousal state prior to stimulation onset significantly influences the effectiveness of auditory stimulation to enhance K-complexes or slow waves in the form of delta power, as well as spindle activity in the form of sigma power. Consequently, pupil size as a proxy for arousal level dynamics may essentially improve current auditory stimulation protocols. Tracking the likelihood of sensory-evoked awakenings upon different sound volumes across pupil level fluctuations and how this relates to the occurrence of slow oscillations may help improve auditory stimulation protocols used in sleep.

Our approach holds the potential to unlock a new research branch within the field of sleep, as it may enable us to track arousal levels during human sleep. This breakthrough allows for fresh fundamental discoveries regarding the interaction between arousal levels and sleep function, potentially offering indirect insight into the LC-NA system during human sleep. Although the current literature in other animal species predominantly supports pupil size to be an indirect readout of LC-mediated arousal during sleep, other mechanisms may be modulating sleep-related pupil size fluctuations directly or indirectly, such as parasympathetically dominant autonomic cholinergic activity, or hypothalamic orexin neurons signaling through the LC^7,17,54^. Moreover, our findings have potential clinical implications in the diagnosis of sleep disturbances. Perturbed arousal levels and LC-function have been closely associated with insomnia, stress-related disorders, and neurodegenerative diseases like Alzheimer’s and Parkinson’s, which often manifest with disrupted sleep patterns^55–58^. By employing our approach, clinicians could directly diagnose whether abnormally elevated arousal levels underlie perturbed sleep in these patients, thus motivating novel treatment approaches.

In conclusion, this study highlights the potential of pupil size as a non-invasive marker of arousal levels during human sleep. Pupil size dynamics correspond to sleep macrostructure and microstructural events, while exhibiting an inverse relationship with spindle ISF. Additionally, the arousal state as indicated by pupil size prior to auditory stimulation influences its effectiveness to enhance delta and sigma power. These findings provide novel insights into the interplay between arousal levels and sleep, opening avenues for research and clinical applications in diagnosing and treating disorders associated with abnormal arousal levels. Pupil size monitoring therefore holds promise as a new tool in both human sleep research and clinical practice.

## Methods

### Participants

In total 18 participants were recruited for this study. One participant was excluded because of predefined exclusion criteria, as they reported an elevated level of eye dryness (score greater than 12 in the Ocular Surface Disease Index questionnaire^59^, 0 to 12 representing normal levels of eye dryness). The 17 included participants (10 female, mean±sd age; 29.83±8.69 years) were non-smokers, reported a regular sleep-wake rhythm, and had a body-mass index between 17-30. Participants reported no presence of psychiatric/neurological diseases, sleep disorders, or clinically significant concomitant diseases. The study was approved by the cantonal Ethics Committee Zurich (reference number: KEKZH, BASEC2022-00340) and conducted in accordance with the declaration of Helsinki. All participants provided written informed consent prior to study participation and received monetary compensation.

### Experiment Procedure

A graphical overview of the experimental procedure is illustrated in Figure 1A. Prior to enrolment, a telephone screening was conducted to explain the procedure and address questions. On the day of testing, participants answered questionnaires about demographics, health status, handedness, sensitivity to noise, eye health, sleep habits, sleep quality, and daytime sleepiness (not reported here). Thereafter, gold electrodes (Genuine Grass electrodes, Natus Medical Inc., Pleasanton, US) were placed to record EEG (Fpz, Cz, O1, O2, M1 (Reference), according to 10-20 system), chin EMG, EOG, and ECG. All channel impedances were kept below 20kΩ. Etymotic insert earphones (Etymotic Research Inc., ER 3C) were then placed in the ear canals and taped to prevent dislodgment during the night. Subsequently, participants laid in a supine position on the bed where the sides of the pillow were wrapped with folded towels to restrict the head from side movements. The right eye remained open by affixing four pieces of tape to the upper eyelid: three strips were placed on the forehead and attached to a fourth strip that was folded outward and positioned onto the upper eyelid. A single, broad strip of tape was used on the lower eyelid (Fig. 1B). To prevent eye dryness, vitamin A eye ointment was applied on the eye immediately before the eye was covered by a transparent eye bandage (PRO Optha S, Lohmann & Rauscher International GmbH & Co. KG, Rengsdorf, Germany). Prior to the application, baby soap was used to clean the plastic dome inside the eye bandage, to minimize the accumulation of water condensation. Finally, the eye tracker goggles (Pupil Core, Pupil Labs GmbH, Berlin, Germany) were placed as shown in Figure 1B. Lights out (approximately 10-11pm) occurred as close as possible to the time participants reported going to bed.

With the exception of the small infrared light source on the goggles, all light sources from the apparatus were covered using black tape or blackout curtains. Once the experimenter turned off the lights and left the sleep lab, participants made 3 up-and-down and 3 left-and-right eye movements that indicated the start of the subject’s sleep window onset of 7.5h. During the complete sleep window, we recorded polysomnography and ECG using a BrainAmp ExG amplifier (BrainProducts GmbH, Gilching, Germany) through OpenVIBE^60^ at a sampling rate of 500 Hz. Additionally, the eye tracker video stream was recorded using the manufacturer’s software, Pupil Capture. The experimenter on call entered the sleep lab up to four hours from taping the eye open to remove the goggles, eye bandage, all tape stripes, and the head restricting towels. Thereafter, the experimenter left the sleep lab again, and the participant was allowed to sleep for the rest of the 7.5h sleep window. In the morning, the experimenter quietly entered the sleep lab, stopped all data acquisition devices, and woke the participant up. The session ended with a second round of sleep quality and mood questionnaires (not reported here) and reimbursement.

### EEG Analysis and Sleep Scoring

All data prior to the moment of the horizontal and vertical eye movements (visually detected in the EOG electrodes) marking the beginning of the sleep window were excluded from further analysis. Then, EEG data was down-sampled to 200Hz using the EEGLAB^61^ toolbox function *pop_resample* in MATLAB (R2019a, MathWorks Inc., Natick, MA). Thereafter, the automatic sleep staging tool YASA^62^ was used to predict sleep stages in 30s windows. Sleep stage predictions with less than 0.6 confidence were visually inspected using the sleep stage visualizer toolbox Visbrain^63^ and manually corrected by an expert scorer following the standard AASM scoring guidelines^33^. Artifacts were rejected based on a semiautomatic artifact removal procedure that was described previously^64^. To determine instantaneous delta and sigma power respectively, Fpz data was bandpass filtered in the 0.5 to 2Hz (delta) frequency range, and Cz data in the 12 to 16Hz (sigma) frequency range using MATLAB 2^nd^ order Butterworth filters. Then, the Hilbert transform of each filtered signal was used to obtain the Hilbert amplitude. Delta and sigma power was thereafter calculated as the square of the absolute Hilbert amplitude.

To estimate the spectral slope of the signal, the Fpz signal was first 0.3Hz high-pass filtered, then 60Hz low-pass filtered, and finally 50Hz notch filtered using a Hamming windowed sinc FIR filter from the EEGLAB toolbox function *pop_eegfiltnew*. The spectral power from 0.5Hz to 45Hz was estimated using a multitaper approach^65^ in the 30s scored epochs which were subsequently used to extract the slope of a fitted model using the FOOOF^66^ algorithm in the 30 to 45Hz range, according to Lendner and colleagues^23^. FOOOF settings were kept default with the exception of minimum peak width limits set to 1 and maximum number of detectable peaks set to 4. Notably, we identified artifacts as 30s windows with power values exceeding 3dB within the frequency range of 0.5 to 45Hz, resulting in the exclusion of 5.45% of the windows.

To get metrics for sleep micro-architecture dynamics (K-complexes, sleep spindles, and arousals), K-complexes present in the Fpz electrode were visually inspected and manually labeled using Visbrain^63^ according to the AASM definition of a K-complex^33^. Due to the inherent difficulty of differentiating K-complexes from increasing background slow wave activity within N3, only K-complexes occurring during N2 and without neighboring slow waves were considered valid. The negative peak of each K-complex was located using MATLAB’s *findpeaks* function. As the short initial positive peak seen in K-complexes is not always present, the onset of the K-complex was defined as 550ms prior to the negative peak as this is typically the time taken for the sharp negative delineation to occur^32,52^. Fast spindles (12 to 16Hz) during N2 were automatically detected using the A7 detector from Lacourse et al.^67,68^. The spindle detection performance was visually inspected for each participant and the sigma correlation parameter for the A7 detector was adjusted accordingly. Spindles lasting less than 300ms or longer than 2s were excluded. Spindle clustering in ‘trains’ was subsequently classified following the Boutin et Doyon^34^ criteria: at least 2 consecutive spindles interspaced by at most 6s. Arousals were automatically detected using an arousal detector algorithm from Fernández-Varela and Alvarez-Estevez^69,70^. The optional “hypnogram” variable was used in the detector algorithm to ensure all detected arousals followed the AASM guidelines^33^. To compare each metric, we excluded epochs where auditory stimuli were presented and normalized pupil size as well as HR to the 5s before each event (Fig. 3).

A cycle-by-cycle approach was applied for the detection of individual sigma power ISFs. First, the Cz sigma power was low-pass filtered at 0.04Hz using a 5th-order Butterworth filter. This cut-off frequency was selected as ISFs have been reported to occur below 0.04Hz in humans^8,9^. Using the MATLAB *findpeaks* function, peaks and troughs were detected. The following criteria had to be met to consider a trough-peak-trough event an ISF: the middle peak and the two troughs occurred during uninterrupted N2; peaks and troughs were at least 15s apart; troughs were within 100s from each other; at least 5% of the fluctuation length contained spindles; no auditory stimulation was presented during the cycle; and no artifacts were present. The ISF analysis depicted in Figure 2 consisted of renormalizing the time scale of sigma power, pupil size, and HR from the time duration of the ISF to the ISF cycle length in terms of percentage cycle. Pupil size and HR were normalized to the average magnitude during each ISF.

### Offline pupil analysis

To calculate pupil size, the eye tracker video was first resampled to 50Hz, and processed using Python 3.8 and DeepLabCut 2.2^21,22^. 36 markers were labeled in each frame as shown in Figure 1B: 12 marking the palpebral fissure circumference, 12 marking the iris circumference, and 12 marking the pupil circumference. Processing of the markers was conducted in MATLAB. First, an ellipse was fitted to the pupil markers. Then, a line was fitted from each iris marker (I) to the centroid of the ellipse (*C*). The intersection point of the fitted line with the ellipse (*P*) was then extracted. Pupil size was calculated as the ratio of the pupil radius to the iris radius using Equation 1, where N is the number of valid iris markers, DIC_i_ is the distance from the i^th^ iris marker to the centroid of the ellipse, and DPC_i_ is the distance from the i^th^ intersection point to the centroid of the ellipse (Fig. 1B).

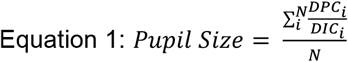

Pupil gaze was calculated by re-referencing the centroid of the pupil markers to the centroid of a polygon generated from the palpebral fissure markers. To account for palpebral fissure twitches (e.g. blink attempts during wake), the palpebral fissure markers were smoothed using a trailing average 40s moving window.

Right eye pupil size was systematically preprocessed using the guidelines and standardized open-source pipeline published by Kret and Sjak-Shie^71^. Invalid pupil diameter samples representing dilation speed outliers and large deviation from trend line pupil size (repeated four times in a multipass approach) were removed using a median absolute deviation (MAD; multiplier in preprocessing set to 12^71^) – an outlier resilient data dispersion metric. Further, temporally isolated samples with a maximum width of 50ms that border a gap larger than 40ms were removed. Data was then linearly interpolated using the MATLAB function *interp1* to match the 200Hz sampling rate of polysomnography data and to fill gaps with up to one second of missing data. Relative pupil size was calculated as a percentage change to the mean pupil size of the last 5s before the events of interest (i.e., spindle trains, K-complexes, arousals, and stimulation windows) or to the average pupil size of the corresponding ISF.

### ECG analysis

ECG R peaks were automatically detected and then visually inspected using the MATLAB-based toolbox PhysioZoo^72^. Data segments consisting of non-detectable peaks or poor quality were excluded from further analyses. HR dynamics is calculated by converting each interval between two R peaks as an inverse of its duration. HR was then linearly interpolated using the MATLAB function *interp1* to match the 200Hz sampling rate of the polysomnography data. Relative HR was calculated as a percentage change to the mean HR of the last 5s before the event of interest (i.e., spindle trains, K-complexes, and arousals) or to the average heart rate of the corresponding ISF.

### Auditory stimulation protocol

During the sleep period, we investigated brain responses to administered auditory stimuli depending on arousal levels. For that, we used our custom-developed EEG-feedback controlled stimulation protocol in OpenViBE^60^ which was an extension of a previously reported protocol by our group^44,48^. As our stimulation condition (verum), we used the 45dB stimulation protocol ISI1 from Huwiler et al.^44^. This consisted of a 1Hz rhythmic EEG feedback-controlled auditory stimulation of a 50ms burst of pink noise with a sound level of 45dB in a windowed 10s ON (auditory stimulation presented) followed by a 20s OFF (no auditory stimulation presented) design. We used different approaches to apply tones for each participant. These included the verum condition (V0, n=4), and a version with the addition of an ISF detection protocol inspired by Cardis et al.^30^ which included the verum condition and a SHAM condition where no auditory stimulation was presented (V1, N=10). The remaining participants were used for the parameter optimization of V1 (V0.5, N=3). For V0 stimulation prerequisites were adapted from our previous work^44^ and for V1 the additional prerequisite of varying sigma power thresholds were implemented to stimulate at different levels of spindle ISF. The analysis in Figure 4A-C included the participants from the V1 protocol. Out of the 10 participants in V1 two participants were excluded in Figure 4A due to having less than 3 stimulations with valid pupil size measurements. This led to a final n=8 for Figure 4A and n=10 for Figures B and C.

To compare the differential effects of high and low arousal levels on responses to auditory stimuli (Fig. 4D-E), we calculated the average pupil size 5s prior to each stimulation window and used it as a baseline (see offline pupil size analysis). Participants with at least 6 stimulation windows with valid pupil data during the 5s baseline were pooled together from V0 (N=2) and V1 (N=5) protocols. Then, we separated each participant into the largest and smallest 25% baseline pupil size ranges for each stimulation condition separately.

### Statistics

Statistical analyses were conducted in R (v 3.6.3; R Core Team, Vienna, Austria). Using the R packages lme4^73^ and lmertest^74^, we computed linear mixed-effects models with several outcome variables (average pupil size, average HR, spectral slope, delta power, or sigma power) and fixed factors with subject as random factor. The fixed factors were sleep stage (Fig. 1F-H), ISF quadrant (Fig. 2G-I), stimulation condition (Fig. 4A-C), or pupil size range and timing of stimulation (Fig. 4D-E). If the one-factor linear mixed-effects models were significant for the fixed effect, we derived post-hoc p-values using Satterthwaite’s method from the R package lmertest and corrected for multiple comparisons with the Hochberg method using the R package emmeans^75^. If the two-factor linear mixed-effect models were significant for the interaction effect (Fig. 5A-B), we derived post-hoc p-values for the contrasts of interest using Satterthwaite’s method from the R package lmertest and corrected for multiple comparisons with the Hochberg method. Visual inspection of the residual plots of the linear models did not reveal any obvious deviations from normality or homoscedasticity. Pupil size (Fig. 3A-C) and HR (Fig. 3D-F) were compared to their respective average values 5s prior to K-complexes, arousals, and arousal using paired t-tests corrected for multiple comparisons using Bejamin-Hochberg correction. To statistically compare verum versus sham conditions (Fig. 4A-C), we averaged the data within three time bins (i) Stim_early_(first 5s of the auditory stimulation), (ii) Stim_late_(last 5s of the auditory stimulation), (iii) Stim_post_(5s bin after auditory stimulation was switched off) and applied paired t-tests between verum and sham for each time bin and were corrected for multiple comparisons with the Benjamini-Hochberg method. To calculate inter-subject correlations of pupil size and spectral slope in 30s epochs across sleep stages, repeated measures correlation analyses were conducted using the R package rmcorr^76^. For the cross-correlation between pupil, HR, and sigma power during ISFs, each signal was zscored within their ISF and then cross-correlated using the MATLAB function *xcorr*. P-values <0.05 were considered significant. Plots were generated using the R package ggplot2^77^ and MATLAB. Where applicable, percentage values reported in the main body of text are in the format mean ± standard error of the mean.

## Acknowledgments

All authors thank their trainees, collaborators, and mentors for the inspiring exchanges on the thematic of this article. The authors would like to express their gratefulness to all participants of the study.

## Supplementary Material

**Supplementary Figure 1.**
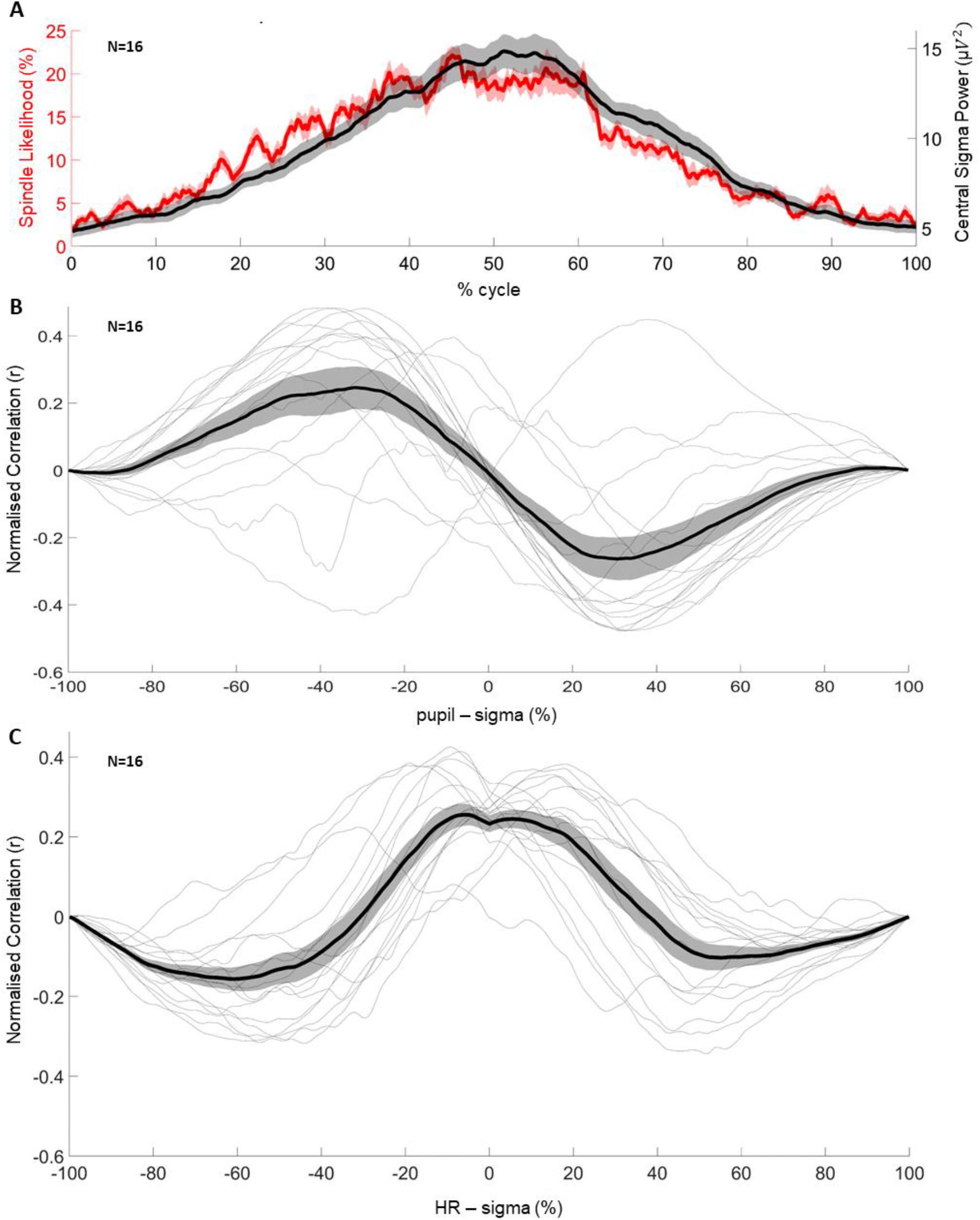
Likelihoods of spindles during infraslow cycles and cross-correlations. (**A**) Spindle likelihood (red & left axis) and sigma power (black & right axis) normalized to the timing of the troughs of detected spindle infraslow fluctuations (ISF). (**B**) Cross-correlation of sigma power (source) and pupil size with the lag being with respect to the cycle percentage of the ISF. (**C**) Cross-correlation of sigma power (source) and heart rate with the lag being with respect to the cycle percentage of the ISF. Centre lines plots represent the mean across participants and shadings around center lines represent the standard error of the mean (SEM).

**Supplementary Figure 2.**
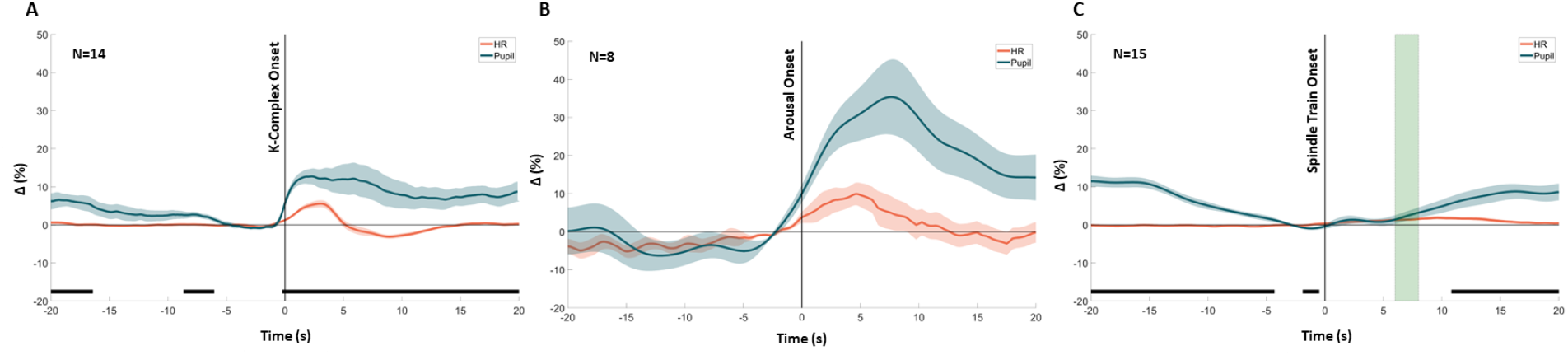
Pupil size compared to instantaneous heart rate dynamics to N2 events. Pupil size color-coded in blue and heart rate color-coded in orange normalized to the 5s prior the detected event: K-complexes (**A**), arousals (**B**), and spindle trains (**C**). Centre lines represent the group mean and shading around the centre lines the SEM. Black bars mark significant (p<0.05) corrected paired t-tests corrected for multiple comparisons (Benjamin-Hochberg correction) between pupil and HR dynamics. Green vertical shading in (**C**) reflects the mean ± 1 SEM of the median spindle train end across participants. (**A**) Significant differences between pupil and HR dynamics surrounding K-complexes are seen during the 20s after its onset and from 5s before up to 20s before. (**B**) No significant differences were found between pupil and HR dynamics surrounding arousals. (**C**) Significant differences between pupil and HR dynamics surrounding spindle trains were present almost everywhere except during the spindle train.

**Supplementary Table 1.** Pupil size contrasts across sleep stages. P-values are those shown in Figure 1F.

| Effect | EMMeans | df | t.ratio | 95% CI (Lower) | 95% CI (Upper) | p-value | Effect size (Cohen's D) |
| --- | --- | --- | --- | --- | --- | --- | --- |
| AWAKE - NREM1 | 0.156 | 60.387 | 10.682 | 0.114 | 0.199 | <0.001 | 3.94 |
| AWAKE - NREM2 | 0.267 | 60.75 | 18.256 | 0.225 | 0.31 | <0.001 | 6.73 |
| AWAKE - NREM3 | 0.326 | 62.898 | 20.921 | 0.281 | 0.372 | <0.001 | 8.23 |
| AWAKE - REM | 0.316 | 64.131 | 19.189 | 0.268 | 0.364 | <0.001 | 7.97 |
| NREM1 - NREM2 | 0.111 | 60.75 | 7.58 | 0.068 | 0.154 | <0.001 | 2.8 |
| NREM1 - NREM3 | 0.17 | 62.898 | 10.91 | 0.125 | 0.216 | <0.001 | 4.29 |
| NREM1 - REM | 0.16 | 64.131 | 9.707 | 0.112 | 0.208 | <0.001 | 4.03 |
| NREM2 - NREM3 | 0.059 | 61.443 | 3.824 | 0.014 | 0.104 | 0.001 | 1.49 |
| NREM2 - REM | 0.049 | 62.641 | 2.994 | 0.001 | 0.097 | 0.008 | 1.24 |
| NREM3 - REM | -0.01 | 60.699 | -0.607 | -0.06 | 0.039 | 0.546 | -0.26 |

**Supplementary Table 2.**
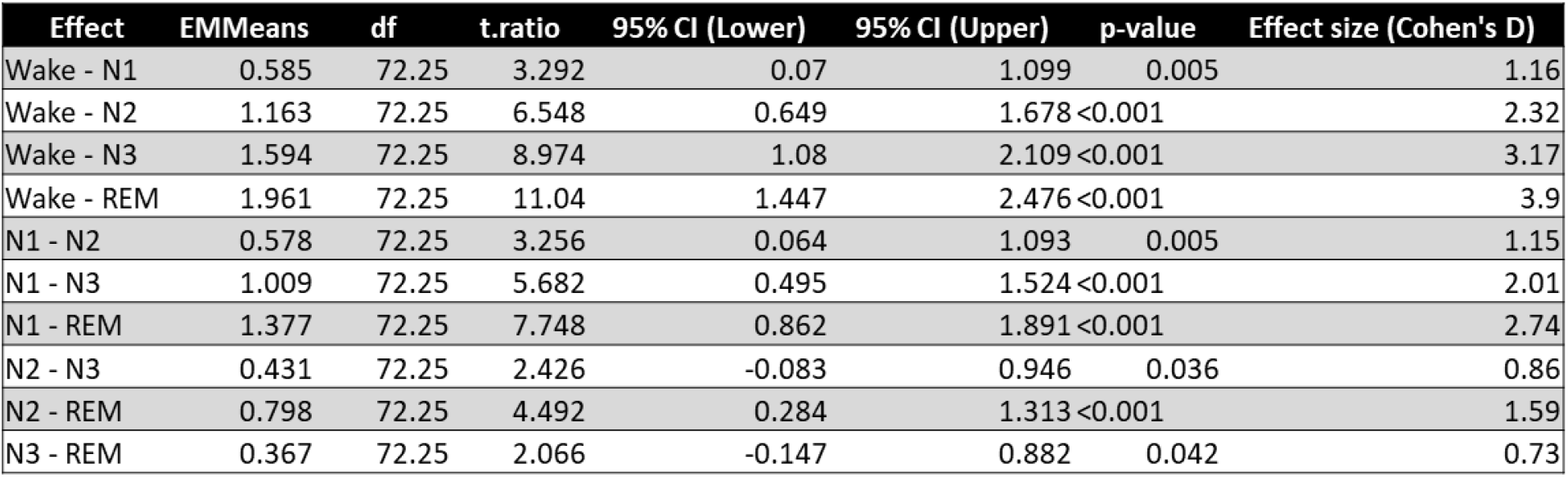
Spectral slope contrasts across sleep stages. P-values are those shown in Figure 1G.

**Supplementary Table 3.** Heart rate contrasts across sleep stages. P-values are those shown in Figure 1H.

| Effect | EMMeans | df | t.ratio | 95% CI (Lower) | 95% CI (Upper) | p-value | Effect size (Cohen's D) |
| --- | --- | --- | --- | --- | --- | --- | --- |
| AWAKE - NREM1 | 5.983 | 59.388 | 4.093 | 1.721 | 10.244 | 0.001 | 1.51 |
| AWAKE - NREM2 | 6.914 | 59.534 | 4.712 | 2.636 | 11.191 | <0.001 | 1.75 |
| AWAKE - NREM3 | 5.323 | 59.837 | 3.376 | 0.728 | 9.919 | 0.01 | 1.35 |
| AWAKE - REM | 2.936 | 59.956 | 1.758 | -1.931 | 7.803 | 0.419 | 0.74 |
| NREM1 - NREM2 | 0.931 | 59.534 | 0.634 | -3.347 | 5.208 | 0.677 | 0.24 |
| NREM1 - NREM3 | -0.66 | 59.837 | -0.419 | -5.255 | 3.936 | 0.677 | -0.17 |
| NREM1 - REM | -3.047 | 59.956 | -1.824 | -7.914 | 1.821 | 0.419 | -0.77 |
| NREM2 - NREM3 | -1.591 | 59.573 | -1.022 | -6.13 | 2.949 | 0.677 | -0.4 |
| NREM2 - REM | -3.977 | 59.695 | -2.412 | -8.784 | 0.829 | 0.133 | -1.01 |
| NREM3 - REM | -2.387 | 59.421 | -1.406 | -7.337 | 2.563 | 0.66 | -0.6 |

**Supplementary Table 4.** Pupil size contrasts across phases of sigma power ISFs. P-values are those shown in Figure 2G.

| Effect | EMMeans | df | t.ratio | 95% CI (Lower) | 95% CI (Upper) | p-value | Effect size (Cohen's D) |
| --- | --- | --- | --- | --- | --- | --- | --- |
| Rise - Peak | -0.011 | 51.2 | -2.271 | -0.024 | 0.002 | 0.058 | -0.83 |
| Rise - Fall | -0.026 | 51.2 | -5.482 | -0.04 | -0.013 | <0.001 | -2 |
| Rise - Trough | -0.011 | 51.2 | -2.247 | -0.024 | 0.002 | 0.058 | -0.82 |
| Peak - Fall | -0.016 | 51.2 | -3.212 | -0.029 | -0.002 | 0.009 | -1.17 |
| Peak - Trough | 0 | 51.2 | 0.024 | -0.013 | 0.013 | 0.981 | 0.01 |
| Fall - Trough | 0.016 | 51.2 | 3.235 | 0.002 | 0.029 | 0.009 | 1.18 |

**Supplementary Table 5.** Heart rate contrasts across phases of sigma power ISFs. P-values are those shown in Figure 2H.

| Effect | EMMeans | df | t.ratio | 95% CI (Lower) | 95% CI (Upper) | p-value | Effect size (Cohen's D) |
| --- | --- | --- | --- | --- | --- | --- | --- |
| Rise - Peak | -0.854 | 51.2 | -3.015 | -1.632 | -0.077 | 0.012 | -1.1 |
| Rise - Fall | -0.106 | 51.2 | -0.376 | -0.884 | 0.671 | 0.709 | -0.14 |
| Rise - Trough | 1.036 | 51.2 | 3.657 | 0.258 | 1.813 | 0.002 | 1.34 |
| Peak - Fall | 0.748 | 51.2 | 2.64 | -0.03 | 1.525 | 0.022 | 0.96 |
| Peak - Trough | 1.89 | 51.2 | 6.672 | 1.112 | 2.667 | <0.001 | 2.44 |
| Fall - Trough | 1.142 | 51.2 | 4.032 | 0.365 | 1.92 | 0.001 | 1.47 |

**Supplementary Table 6.** K-complex likelihood contrasts across phases of sigma power ISFs. P-values are those shown in Figure 2I.

| Effect | EMMeans | df | t.ratio | 95% CI (Lower) | 95% CI (Upper) | p-value | Effect size (Cohen's D) |
| --- | --- | --- | --- | --- | --- | --- | --- |
| Rise - Peak | -2.629 | 51.2 | -5.599 | -3.917 | -1.34 | <0.001 | -2.04 |
| Rise - Fall | -2.225 | 51.2 | -4.74 | -3.514 | -0.937 | <0.001 | -1.73 |
| Rise - Trough | -0.228 | 51.2 | -0.486 | -1.517 | 1.06 | 0.629 | -0.18 |
| Peak - Fall | 0.403 | 51.2 | 0.859 | -0.885 | 1.692 | 0.629 | 0.31 |
| Peak - Trough | 2.4 | 51.2 | 5.112 | 1.112 | 3.689 | <0.001 | 1.87 |
| Fall - Trough | 1.997 | 51.2 | 4.254 | 0.708 | 3.286 | <0.001 | 1.55 |

**Supplementary Table 7.** Pupil size and heart rate during the 5s prior to N2 events versus general N2 pupil size.

| Metric | Effect | Mean % change | S.E.M % change | t | df | p-value |
| --- | --- | --- | --- | --- | --- | --- |
| Pupil size | K-complex - N2 | -8.30 | 2.31 | -2.174 | 41.9 | 0.071 |
|  | Arousals- N2 | 0.95 | 4.94 | 0.365 | 43.3 | 0.071 |
|  | Spindles - N2 | -8.34 | 2.75 | -2.297 | 41.5 | 0.717 |
| Heart rate | K-complex - N2 | -1.03 | 0.46 | -1.091 | 43.2 | 0.382 |
|  | Arousals- N2 | 3.50 | 1.59 | 3.202 | 43.4 | 0.008 |
|  | Spindles - N2 | -0.89 | 0.76 | -0.884 | 43.2 | 0.382 |

